# Glutaredoxin 3 (GLRX3) confers a fusion oncogene-dependent vulnerability to Ewing sarcoma

**DOI:** 10.1101/2024.04.24.590877

**Authors:** Endrit Vinca, Anna C. Ehlers, Alina Ritter, David Obermeier, Cornelius M. Funk, Florian H. Geyer, Melissa Schmucker, Jing Li, Malenka Zimmermann, A. Katharina Ceranski, Fabia Fuchslocher, Christina Mertens, Ruiyue Qiu, Martina M. Muckenthaler, Alina Dahlhaus, Silvia von Karstedt, Roland Imle, Ana Banito, Javier Alonso, Heike Peterziel, Olaf Witt, Ina Oehme, Florencia Cidre-Aranaz, Thomas G. P. Grünewald, Shunya Ohmura

## Abstract

Ewing sarcoma (EwS) is a highly aggressive bone and soft-tissue associated cancer for which there are no effective targeted therapeutics available. Genetically, EwS is driven by aberrantly active EWSR1::ETS fusion transcription factors, most commonly EWSR1::FLI1. Despite their unique expression in EwS, all attempts to effectively target these fusion oncoproteins clinically were not yet successful, wherefore alternative targets are required.

Here, we functionally characterize the evolutionarily conserved oxidative stress regulator glutaredoxin 3 (GLRX3) as a EwS-specific and EWSR1::FLI1-dependent vulnerability. Through integration of transcriptome-profiling, conditional drug screens in 3D cultures, and functional experiments, we discover that GLRX3 promotes EwS growth in vitro and in vivo, and that it has a key role in mitigation of oxidative stress and maintenance of iron homeostasis. These GLRX3 functions can be exploited in both GLRX3-high and -low expressing EwS cells by targeted therapeutics including CDK4/6 inhibitors and inducers of apoptotic and ferroptotic cell death. Collectively, our results exemplify how the interplay of an evolutionarily conserved oxidative stress regulator with a dominant oncogene can promote malignancy but provide opportunities for predictive diagnostics and personalized therapy.

## INTRODUCTION

Ewing sarcoma (EwS) is the second most common malignant bone and soft-tissue cancer in children, adolescents, and young adults^1^. Characterized by a highly undifferentiated embryonal phenotype, EwS poses a clinical challenge as a rapidly metastasizing cancer, with ∼25 % of cases presenting with metastasis at diagnosis^1,2^. Despite the application of highly toxic and in part mutilating therapies, approximately one-third of patients still succumb to the disease^1,2^. To date, specific targeted therapeutics for EwS are not available^1,2^. Thus, there is an urgent need for novel, more effective, and less toxic therapeutic options.

Genetically, EwS is characterized by chromosomal translocations leading to the expression of pathognomonic *FET::ETS* gene fusions with *EWSR1::FLI1* being the most common one (∼85 % of cases,)^3,4^. The encoded FET::ETS fusion proteins act as potent oncogenic transcription factors that drive the malignant and highly aggressive phenotype of EwS^1^. Since FET::ETS fusions are specific to EwS cells, and as EwS cells are addicted to the oncogenic FET::ETS activity, multiple efforts have been undertaken to specifically target and neutralize these fusions^5–8^. However, it has been shown that complete suppression of their oncogenic activity may not be feasible^1,5,9^, and that incomplete inhibition may render EwS cell even more aggressive^10,11^. Indeed, due to their lack of enzymatic activity, intranuclear localization, in part unstructured protein conformation, the ubiquitous expression of their constituting wildtype proteins, and generally low immunogenicity, FET::ETS fusion oncoproteins have been largely classified as ‘undruggable’^5,9,12^.

To identify alternative targets, several comprehensive screening efforts, such as The Cancer Dependency Map Project (DepMap) of the Broad Institute, have been undertaken to systematically map cancer vulnerabilities across tumor entities including EwS^13^. In fact, DepMap has highlighted several potential targets for EwS. Some of those are currently being followed-up mechanistically^14–17^, but for many other candidates functional validation is still lacking.

In this report, we leveraged existing DepMap data and combined them with original functional experiments to identify targeting of the multifunctional protein glutaredoxin 3 (GLRX3) as a preferential vulnerability for EwS cells depending on the expression of EWSR1::FLI1 fusions. Using a functional genomics approach, we demonstrate that GLRX3 confers a pro-proliferative phenotype on EwS cells in vitro, in vivo, and in situ, that is related to its function in redox regulation and cellular iron homeostasis. Finally, we provide evidence that GLRX3 on the one hand protects EwS cells from induction of apoptotic and ferroptotic cell death mediated by the small molecule drugs, and on the other hand that GLRX3 sensitizes EwS cells toward CDK4/6 inhibitors, which may have important translational implications for the design of personalized treatment strategies.

## RESULTS

### EWSR1::FLI1 confers a preferential vulnerability toward GLRX3 inhibition in EwS

To identify candidate target structures for therapeutic intervention with preference for EwS cells, we explored curated DepMap data^18,19^. To this end, we calculated the Δ gene effect for EwS by subtracting the average gene effect in non-EwS cell lines (*n*=1,077) for a given gene from the average gene effect in EwS cell lines (*n*=23). In a second step, we calculated the individual significance levels for each Δ gene effect and plotted the resulting adjusted *P*-values against the actual Δ gene effects in EwS (**Fig. 1a**). As shown in **Fig. 1a**, this analysis yielded many previously identified and described EwS-specific vulnerabilities, such as *ETV6*, *STAG1*, and *TRIM8*^14–17,20^ as well as *EWSR1* and *FLI1* – the constituents of the EwS-specific *EWSR1::FLI1* fusion – as top hits. Of note, we also identified the new candidate gene *glutaredoxin 3* (*GLXR3*), which is an evolutionary conserved multifunctional protein with implications in redox regulation and iron/sulfur-homeostasis^21–23^. Iron-sulfur clusters (ISC) are trafficked within the cytosol by GLRX3 (ref. ^24^). They play a crucial role in the maturation of numerous metalloproteins and are well known for their role in electron transport in mitochondria via redox reactions^24^. It has been described, that ISC trafficking by GLRX3 may influence the cellular function of the iron-responsive element-binding protein 1 (IRP1) – a bifunctional protein also known as aconitase (ACO1)^25^. Aconitases require 4Fe-4S-clusters for their enzymatic activity, consequently they exhibit an enzymatic role in an iron saturated cell state, but function as an iron regulatory protein in an iron-depleted cell state by binding to iron responsive elements (IRE) and regulating the expression of transferrin-receptor 1 (TfR1) and ferritin, increasing iron uptake and storage^26,27^. Additionally, ISC are required in DNA polymerases and DNA helicases to function properly and maintain genomic stability^28–32^. Accordingly, depletion of GLRX3 has been associated with altered cellular redox homeostasis in nasopharyngeal carcinoma and immortalized T-lymphocytes, resulting in increased oxidative stress levels^22,23^. This finding was of particular interest to us because we previously reported on a delicate regulation of redox homeostasis in EwS, which contributes to its aggressiveness^33,34^. In line with these findings, we found that *GLRX3* ranks among the top EwS-specific gene dependencies when ranking the Δ gene effect of *GLRX3* for EwS compared to other cancer entities included in DepMap (**Fig. 1b,c**). Since *EWSR1::ETS* fusions are pathognomonic for EwS, we hypothesized that this EwS-specific gene dependency on *GLRX3* may be functionally related to the presence of these gene fusions. To test this hypothesis, we employed a heterologous model derived from cervical cancer cells (HeLa) that harbor doxycycline (Dox)-inducible expression system for either wildtype *FLI1* or *EWSR1::FLI1* as previously described^35^. Using these genetically modified HeLa cells, we tested whether the RNA interference-mediated knockdown of *GLRX3*, which is expressed in HeLa cells at levels comparable to EwS cell lines (**Supp. Fig. 1a**), results in changes of viable cell counts depending of ectopic expression of either *FLI1* or *EWSR1::FLI1*. Strikingly, as displayed in **Fig. 1d**, knockdown of *GLXR3* onto ∼5 % remaining expression (**Supp. Fig. 1b**) exclusively affected cell viability upon concomitant induction of *EWSR1::FLI1*, but not upon induction of *FLI1* or under control conditions (Dox –). Similarly, transient knockdown of *GLXR3* in two EwS cell lines that contained either a dox-inducible shRNA against *EWSR1::FLI1* (A-673/TR/shEF1 and MHH-ES-1/TR/shEF1)^36^ led to a significant reduction of viable cell counts 96 h after transfection only in the presence of the fusion (Dox –), while this effect was abrogated upon fusion knockdown (Dox +) (**Fig. 1e**). Collectively, these results indicated that *GLRX3* constitutes a preferential vulnerability for EwS cell lines that is conferred by the presence of *EWSR1::FLI1*.

**Fig. 1.**
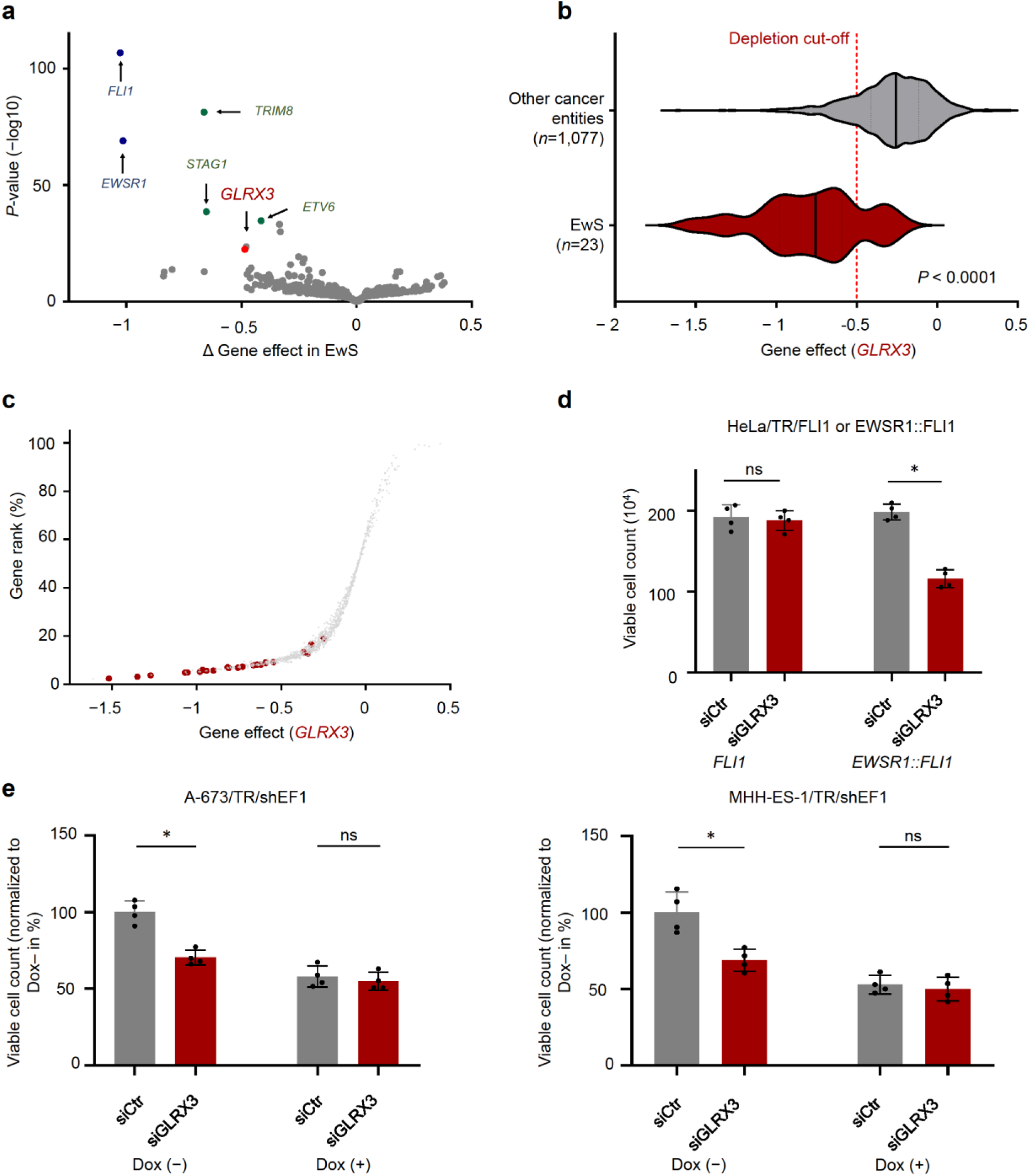
EWSR1::FLI1 confers a preferential vulnerability toward GLRX3 inhibition in EwS. **a)** Volcano plot for Δ gene effects plotted against individual statistical significance values for each Δ gene effects in comparison between EwS and non-EwS cell lines. Δ gene effects: (average gene effects of a given gene in EwS) – (average gene effects of the gene in non-EwS cell lines). **b)** Comparison of the *GLRX3* gene effect in 23 EwS cell lines with 1,077 other cancer cell lines. Vertical bars represent means and dotted lines quartiles. A score less than –0.5 represents ‘depletion’, while a score of –1 is comparable to the median of all pan-essential genes. *P*-value was calculated via two-tailed unpaired t-test with Welch’s correction. **c)** Scatter plot for *GLRX3* gene effects in all cell lines included in DepMap and normalized *GLRX3* dependency rank. Each dot indicates one cell line and EwS are highlighted by red color. **d)** Cell viability analysis of either FLI1 or EWSR1::FLI1 induced HeLa cells upon transient transfection of either non-targeting siRNA or siRNA against *GLRX3*. Viable cells were quantified by using trypan blue dye exclusion method. *n*=4 biologically independent experiments. Horizontal bars represent means and whiskers SEM. *P*-values determined via two-tailed Mann-Whitney test. **e)** Cell viability analysis of EwS cell lines A-673 and MHH-ES-1 with EWSR1::FLI1 high (Dox–) or low (Dox+) upon transient transfection of either non-targeting siRNA or siRNA against *GLRX3*. Viable cells were quantified by using trypan blue dye exclusion method. *n*=4 biologically independent experiments. Horizontal bars represent means and whiskers SEM. *P-*values determined via two-tailed Mann-Whitney test. \**P* < 0.05.

### GLRX3 promotes proliferation and tumorigenicity in EwS

As GLRX3 is a multifunctional protein with complex roles depending on the given cell type^23,25,37–42^, we sought to obtain first insights into its potential roles in EwS by investigating available gene expression data derived from EwS patient tumors. To that end, we employed our large patient cohort (*n*=196) with matched gene expression and clinical data^43^ and computed the Pearson correlation for gene expression of all genes of this dataset against *GLRX3*. Ranking all genes by their Pearson correlation coefficient and conducting a fast gene set enrichment analysis (fGSEA) on these data suggested that *GLRX3* in EwS is associated with enrichment of gene signatures involved in cell proliferation, cell cycle regulation, and DNA replication (**Fig. 2a**). To functionally validate these bioinformatic predictions, we transduced three EwS cell lines (A-673, EW-22, and MHH-ES-1) that exhibited *GLRX3* levels representative for the majority of EwS cell lines (**Supp. Fig. 1c**) with the pLKO-TET-ON vector enabling Dox-inducible expression of two different specific shRNAs against *GLRX3* or a non-targeting control shRNA. As shown in **Supp. Fig. 1d**, Dox-treatment of these cell lines led to reduced *GLRX3* expression to 6–43% compared to the non-targeting control shRNA and Dox– conditions. In keeping with the patient-derived data (**Fig. 2a**), subsequent functional analysis demonstrated that *GLRX3* silencing is accompanied by a strong reduction of viable cell counts across shRNAs and cell lines (**Fig. 2b**) in short-term (96 h) proliferation assays, and a remarkably reduced capacity of clonogenic cell growth upon long-term knockdown (11 d) (**Fig. 2c**). To investigate whether these phenotypes may be attributed to slowdown of cell cycle or concomitant increases in cell death, we carried out functional assays. As shown in **Fig. 2d**, knockdown of *GLRX3* had a consistent and strong effect on G1/S-transitions during cell cycle across cell lines, which was accompanied in some cell lines by a variable extent of subsequent cell death as indicated by Annexin V/PI-stains as well as cell counting assays using the trypan blue exclusion method (**Supp. Figs. 2a,b**). To confirm these in vitro results, we carried out subcutaneous xenotransplantation experiments in NSG mice with/without Dox-inducible knockdown of *GLRX3* once tumors were palpable. Strikingly, conditional silencing of *GLRX3* led to arrest of tumor growth and in some instances even tumor regression in two independent cell lines (A-673 and MHH-ES-1) – an effect not observed in non-targeting control shRNA xenografts (**Fig. 2e**). In accordance with the in vitro results, (immuno)histological analyses of the xenografts demonstrated a marked reduction of mitotic cell counts upon *GLRX3* knockdown (**Fig. 2f**). In sum, these in situ, in vitro, and in vivo results demonstrated that *GLRX3* is critical for EwS proliferation and tumorigenesis.

**Fig. 2.**
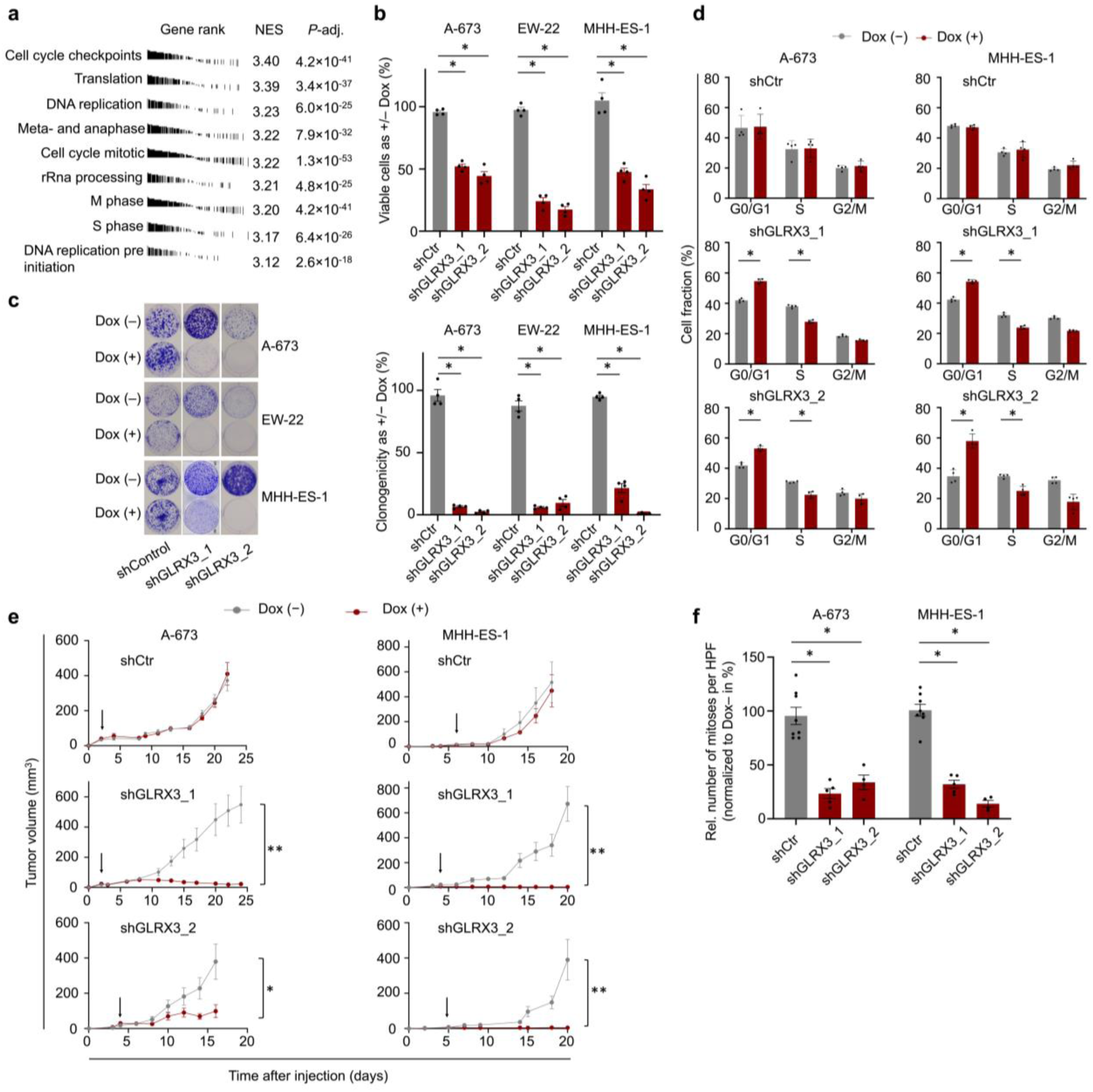
GLRX3 promotes proliferation and tumorigenicity in EwS. **a)** Gene set enrichment analysis (GSEA) for *GLRX3* co-expressed genes derived from gene expression data sets of 196 EwS tumor samples. Genes were pre-ranked according to Pearson correlation coefficients between *GLRX3* and other genes and enrichment scores for each gene set were calculated by GSEAPreranked module. NES, normalized enrichment score. padj, adjusted *P-*value. **b)** Cell viability analysis of EwS cell lines A-673, EW-22, and MHH-ES-1 upon induction of either non-targeting shRNA (shCtr) or two different specific shRNA against *GLRX3* (shGLRX3). Viable cells were quantified by using trypan blue dye exclusion method and values were normalized to those of Dox–. *n*=4 biologically independent experiments. Horizontal bars represent means and whiskers SEM. *P-*values determined via two-tailed Mann-Whitney test. **c)** Left: representative images and right: analysis of clonogenic growth of EwS cell lines A-673, EW-22, and MHH-ES-1 upon induction of either non-targeting shRNA (shCtr) or two different specific shRNA against *GLRX3* (shGLRX3). Colony intensity was analyzed by using ImageJ plugin ColonyArea^74^ and values were normalized to those of shCtr. *n*=4 biologically independent experiments. Horizontal bars represent means and whiskers SEM *P*-values determined via two-tailed Mann-Whitney test. **d)** Cell cycle analysis of EwS cell lines A-673 and MHH-ES-1 upon induction of either non-targeting shRNA (shCtr) or two different specific shRNA against *GLRX3* (shGLRX3). EwS cells stained with propidium iodide (PI) were detected by flow cytometry and cellular fractions for each cell cycle phase was analyzed using FlowJo. **e)** Tumor growth analysis of EwS cell lines A-673 and MHH-ES-1 harboring two different Dox-inducible shGLRX3 (shGLRX3_1 or shGLRX3_2) or non-targeting shRNA (shCtr) xenografted in NSG mice. Once tumors were palpable, animals were randomized in Dox (+) (*n*=8) or Dox (–) (*n*=8) group and tumor diameters were measured by a caliper. Arrows indicate the time point of start of Dox-treatment. Two-sided Mann-Whitney test. **f)** Mitoses of xenografts described in (e) were quantified in HE-stained slides. *n*≥4 samples per condition. Horizontal bars represent means, and whiskers represent the SEM. *P*-values determined via two-tailed Mann-Whitney test. \*\**P* < 0.01, \**P* < 0.05.

### GLXR3 silencing is associated with dysregulation of oxidative stress levels and cellular iron homeostasis in EwS

To gain deeper insights into the underlying molecular changes induced by GLRX3 in EwS cells that may account for the observed phenotypes (**Figs. 1** and **2**), we carried out transcriptome profiling of two EwS cell lines (A-673 and EW-22) 96 h after initiation of Dox-inducible shRNA-mediated knockdown of *GLRX3*. Since we profiled the same EwS cell lines with/without knockdown of EWSR1::FLI1 for 96 h in frame of our previous Ewing Sarcoma Cell Line Atlas (ESCLA) project on the same Affymetrix DNA microarrays (human Clariom D)^36^, we first generated a harmonized dataset of our new and former microarrays by preprocessing and normalizing the data jointly (see methods). Due to the strong dependency of EwS cells on *GLRX3* that is apparently mediated by the co-expression of EWSR1::FLI1 (**Figs. 1d,e**), we examined the potential overlap of up- or downregulated differentially expressed genes (DEGs) in both conditions. As shown in **Fig. 3a**, EWSR1::FLI1 and GLRX3 share a statistically significant (*P*<0.0001) proportion of commonly up- or downregulated DEGs. Subsequent overrepresentation analyses using PANTHER revealed that the commonly upregulated genes were mainly enriched in molecular signatures involved in cell proliferation and cell cycle regulation (**Fig. 3b, Supp. Table 1**), which is consistent with our observation made from our in vitro and in vivo models as well as the patient cohort (**Fig. 2**).

**Fig. 3.**
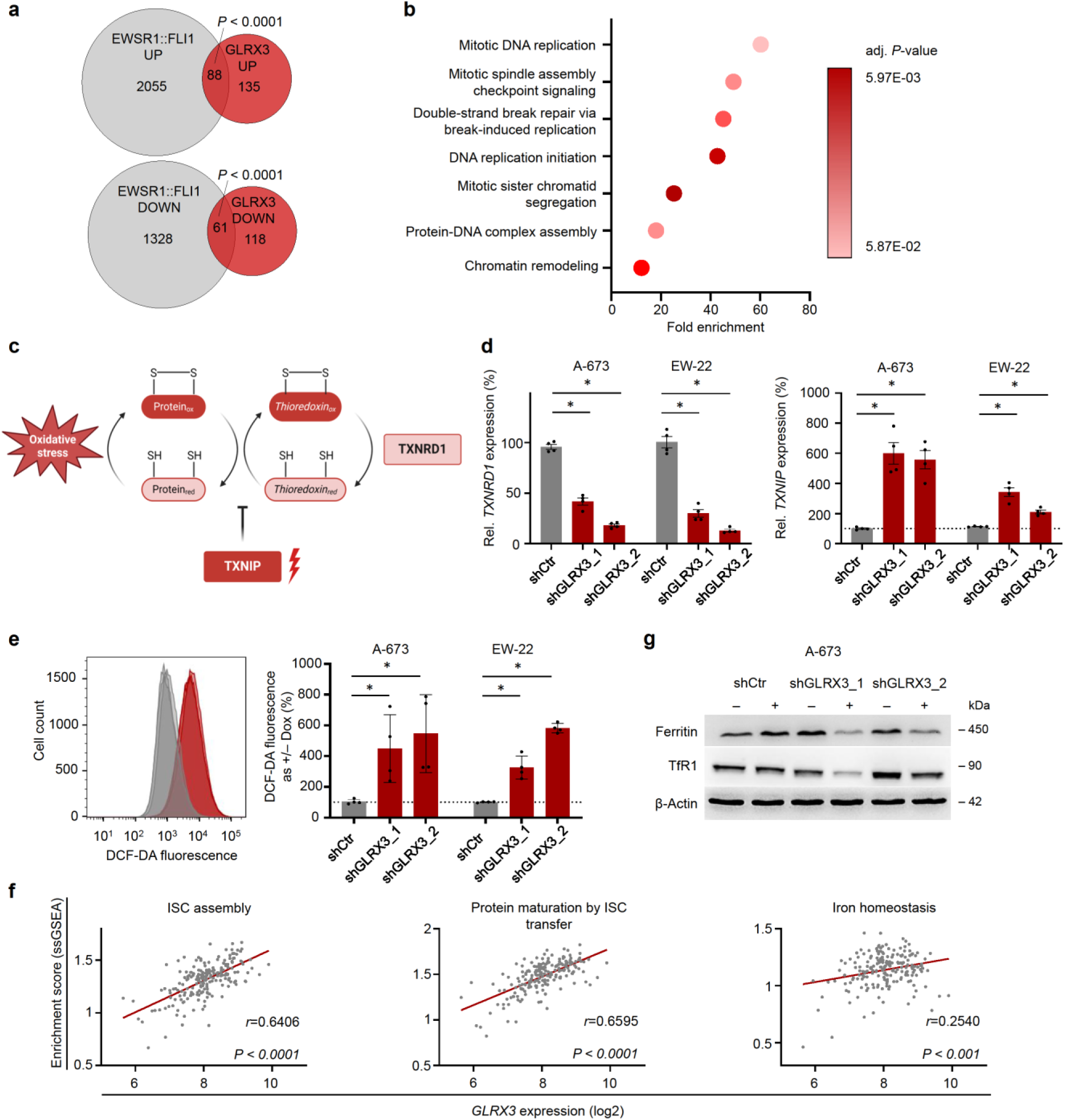
GLXR3 silencing is associated with dysregulation of oxidative stress levels and cellular iron homeostasis in EwS. **a)** Overlap of up- and downregulated DEGs between *EWSR1::FLI1* high/low and *GLRX3* high/low transcriptome in A-673 and EW-22. DEGs were filtered for adj. *P*-value < 0.05 and |logFC| > 1. **b)** GO overrepresentation analysis of commonly upregulated DEGs between *EWSR1::FLI1* high/low and *GLRX3* high/low using PANTHER to classify gene according to GO term PANTHER-Slim Biological Process, filtering through an adj. *P*-value >0.05. The fold enrichments display the genes observed in our DEG list over the expected number of genes of a reference list. If the fold enrichment is >1, the category is overrepresented. **c)** Schematic illustration of the effects of *TXNRD1* and *TXNIP* on the thioredoxin pathway. **d)** Left: *TXNRD1* expression. Right: *TXNIP* expression. *n*=4 biologically independent experiments. Horizontal bars represent means and whiskers SEM. *P*-values determined via two-tailed Mann-Whitney test. **e)** Correlation analysis between *GLRX3* expression and the Enrichment Score (ES) assessed by ssGSEA performed on gene expression data sets of 196 EwS tumors. Lines indicate linear regressions. *P-*values determined by two-sided Pearson correlation test. **f)** Representative histogram (left) and quantification (right) of DCF-DA fluorescence in A-673 and EW-22 cells after 96 h *GLRX3* silencing compared to shCtr. Horizontal bars represent means, *n*=4 biologically independent experiments. *P*-values determined via two-sided Mann-Whitney test. **g)** Representative ferritin and transferrin-receptor (TfR1) western blot of *n*=3 biologically independent replicates. \**P* < 0.05.

As *GLRX3* is a multifaceted protein with diverse cellular roles including redox and iron homeostasis by ISC trafficking^44,45^, we next scrutinized our list of DEGs that were exclusively regulated by GLRX3 but not EWSR1::FLI1 specifically for genes involved in these processes. Strikingly, the most upregulated genes after *GLRX3* silencing (∼20-fold upregulation, *P*=0.001) was *thioredoxin inhibiting protein* (*TXNIP*) – a protein that we identified previously to be involved in redox homeostasis of EwS^33^ – while among the most downregulated DEGs, we found *thioredoxin reductase 1* (*TXNRD1*) as the 4^th^ most downregulated gene (∼7-fold downregulation, *P*=0.013) (**Supp. Table 2**). Physiologically TXNRD1 buffers cellular oxidative stress, and its activity is tightly controlled by TXNIP (**Fig. 3c**)^46–48^. Since the antagonistic regulation of both genes strongly suggested an involvement of GLRX3 in redox homeostasis, we first confirmed their strong dysregulation upon GLRX3 knockdown in independent experiments by qRT-PCR (**Fig. 3d**). In a second step, we validated changes in cellular oxidative stress levels by subjecting EwS cells with/without *GLRX3* silencing to dichlorodihydrofluorescein diacetate (DCF-DA) to dichlorodihydrofluorescein (DCF) conversion assays. As shown in **Fig. 3e**, flow-cytometric assessment of DCF-DA to DCF conversion demonstrated that silencing of *GLRX3* led to a ∼5-fold increase of oxidative stress levels in EwS cells.

Previous studies have suggested, that ISC bound to GLRX3 could serve as a cellular sensor for elevation of oxidative stress levels by dissociation of these clusters after redox-induction^23,25,44^. We were able to validate the potential link between GLRX3 and ISC assembly, protein maturation by ISC transfer and iron homeostasis in EwS patient tumors by correlating the enrichment scores of these pathways, calculated by ssGSEA, with *GLRX3* expression (**Fig. 3f**). Consistent with these molecular gene signatures, we found that *GLRX3* silencing for 96 h led to downregulation of ferritin as evidenced by western blotting, while for the transferrin receptor the effect was less pronounced (**Fig. 3g**). In sum, these results indicated that GLXR3 is involved in regulation of oxidative stress levels and cellular iron homeostasis in EwS.

### GLRX3 may serve as a predictive biomarker for targeted therapy of EwS patients

As our results indicated that *GLRX3* may serve as a selective vulnerability in EwS, we sought to therapeutically target this protein. Unfortunately, to the best of our knowledge, there is currently no specific targeted drug available for GLRX3, wherefore we asked whether we could use GLRX3 as predictive biomarker for other targeted therapeutics that interfere with GLRX3 associated pathways. Indeed, as shown in **Supp. Fig. 2c**, *GLRX3* is heterogeneously expressed across EwS tumors in our patient cohort, wherefore we tested whether we could identify potential inroads for targeted intervention for both *GLRX3*-high and -low patients. To this end, we followed an orthogonal strategy by combining an agnostic drug screen on EwS cell lines with/without *GLRX3* silencing with a hypothesis-driven approach based on our original findings (**Figs. 1–3**) and prior data in the literature on the molecular function of this protein.

For the drug screen, A-673 cells with a Dox-inducible shRNA against *GLRX3* were grown in 3D spheres and pre-treated with Dox for 48 h and subsequently exposed to 87 different clinically relevant drugs in ascending concentrations for another 72 h. Subsequent read-out using CellTiter-Glo yielded dose-response curves from which we calculated Δ drug sensitivity scores (ΔDSS) in both GLRX3-high (Dox–) and GLRX3-low (Dox+) conditions as previously described^49^ (**Fig. 4a**). Of the 87 drugs, 9 had a Δ DSS ≥ 5 and 6 had a Δ DSS ≤ –5. Among the drugs with higher efficacy in GLRX3-high conditions, we focused in independent validation experiments – using a colorimetric resazurin assay^50^ – on the CDK4/6 inhibitor palbociclib because it is currently in clinical trials that included EwS patients^51^ (*NCT04129151*, *NCT03155620*, *NCT03526250*, *NCT03709680*) and as we and others have shown previously that CDK4/6 inhibition is effective in preclinical EwS models^52–54^. Furthermore, we also paid particular attention to CDK4/6 inhibitors because the mechanism is based on a disruption of the cell cycle machinery, and since our data pointed to a strong connection of GLRX3 and cell cycle signatures in vitro and in situ (**Fig. 2**). In parallel, we assessed the response of EwS cells to ribociclib – another FDA-approved CDK4/6-inhibitor – as an additional control. As shown in **Fig. 4b**, silencing of *GLRX3* significantly increased the IC50 values for both CDK4/6 inhibitors (∼2–5.6-fold) in two cell lines (palbociclib: A-673 [7.9 µM → 18.1 µM], MHH-ES-1 [3.8 µM → 11.3 µM]; ribociclib: A-673 [12.2 µM → 41.1 µM], MHH-ES-1 [6.7 µM → 34.5 µM]). Among the most effective drugs in GLRX3-low conditions was the pro-apoptotic drug navitoclax, which is a BCL-2 inhibitor currently in clinical trials for several oncological indications (*NCT00445198*, *NCT01989585*, *NCT06210750*, *NCT02079740*, *NCT05740449*). Strikingly, these results could be fully validated in independent experiments using colorimetric resazurin drug response assays in both cell lines and across shRNAs with a differential response of ∼38-fold (A-673 [19.5 µM → 0.4 µM], MHH-ES-1 [8.4 µM → 0.3 µM]) (**Fig. 4c**), suggesting a relative resistance of GLXR3-high cells toward this drug.

**Fig. 4.**
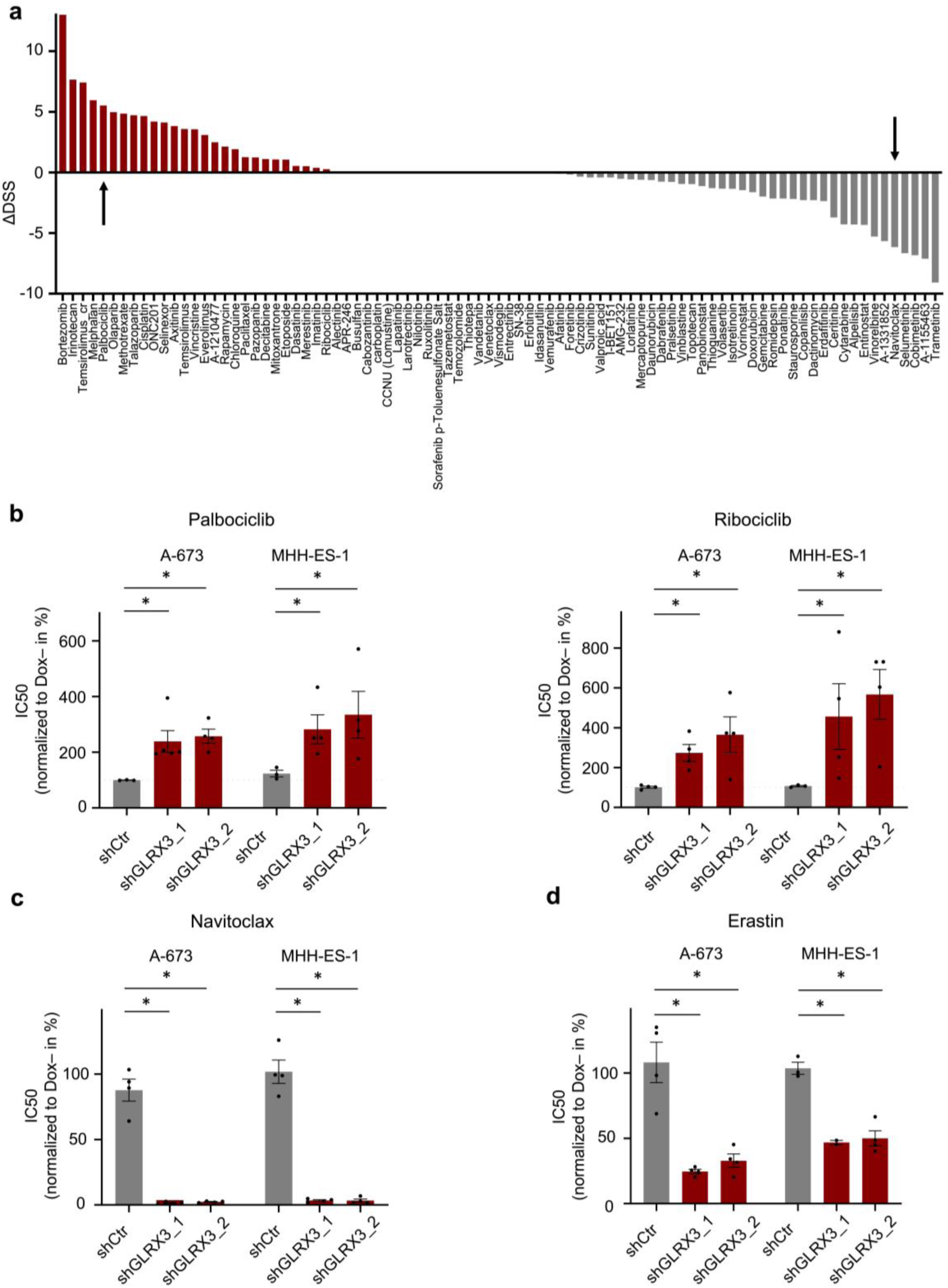
GLRX3 as a potential predictive biomarker for targeted therapy of EwS. **a)** Waterfall plot for differential drug sensitivity in A-673/TR/shGLRX3 with/without *GLRX3* silencing. For each condition drug sensitivity scores (DSS) were determined by the extended DSS^49,75^ based on the integral of an advanced five-parameter dose-response curve model^49^. ΔDSS: (DSS for without *GLRX3* silencing) – (DSS for with *GLRX3* silencing). ΔDSS ≥5 indicates higher sensitivity for high *GLRX3* condition. Drugs selected for functional validation are indicated with black arrows. **b-d)** Drug sensitivity analysis for (b) palbociclib/ribociclib, (c) navitoclax, and (d) erastin in EwS cell lines A-673 and MHH-ES-1 upon induction of either non-targeting shRNA (shCtr) or two different specific shRNA against *GLRX3* (shGLRX3). IC50 values determined by Resazurin assays were normalized to those for without *GLRX3* silencing. *n≥*4 biologically independent experiments. Horizontal bars represent means and whiskers SEM. *P*-values determined via two-sided Mann-Whitney test. \*\**P*<0.01, \**P* < 0.05.

Since the conclusions drawn from both the agnostic drug-screens as well as the hypothesis-driven investigation on the proliferative phenotype, conferred by GLRX3, support the use of cell cycle inhibitors, such as CDK4/6 inhibitors, in GLRX3-high EwS tumors, we wondered whether we could therapeutically exploit the potential dysregulation of cellular iron-homeostasis seen in GLRX3-low EwS cells (**Fig. 3**). Since changes in cellular iron-homeostasis and oxidative stress regulation have been shown to be major contributors for the induction of iron-dependent cell death (i.e. ferroptosis)^55–57^, we hypothesized that GLRX3 may serve as a predictive biomarker for ferroptosis-inducers, such as erastin^55,58,59^. To test this hypothesis, we conducted drug-response assays using erastin in two EwS cell lines (A-673, MHH-ES-1) with/without *GLRX3* silencing. In keeping with our hypothesis, we observed a ∼4-fold decrease of the IC50 values in GLRX3-low cells (**Fig. 4d**). In synopsis, these results suggest that GLRX3 may serve as a predictive biomarker for multiple drugs that are related to cell cycle inhibition and/or apoptotic/ferroptotic cell death.

## DISCUSSION

In this study, we show that the multifaceted protein GLRX3 constitutes a preferential vulnerability for *EWSR1::FLI1*-driven EwS cells, promoting cell proliferation and tumor growth. We demonstrate that GLRX3 is implicated in oxidative stress and cellular iron-metabolism, and that its variable expression in EwS tumors may have relevance as a predictive biomarker for a diverse array of drugs related to cell cycle inhibition and/or apoptotic/ferroptotic cell death.

By analysis of curated DepMap data in conjunction with functional experiments, we validate that *GLRX3* silencing is deleterious for EwS cells – a phenotype that appears to be mediated by coexpression of EWSR1::FLI1 fusion oncoproteins as evidenced in a variety of EwS cell lines as well as in heterologous models (**Fig. 1d,e**). In support of this finding, preliminary comparative transcriptome profiling experiments show that EWSR1::FLI1 and GLRX3 share common sets of DEGs in EwS cells, such as those involved in cell cycle and proliferation, as well as canonical EWSR1::FLI1 targets (**Supp. Fig. 2d, Supp. Table 3**), which may underpin the observed dependency on GLRX3 conditional to the fusion. Yet, the precise mechanism of this delicate interaction needs to be elucidated in future studies. While this dependency appears to depend in EwS models on EWSR1::FLI1 fusions, other studies in oral squamous cell carcinoma and pancreatic adenocarcinoma showed an albeit less pronounced dependency of the corresponding models^60–62^, which may hint to the fact that also other driver oncogenes could mediate such a dependency and that this phenomenon is not exclusive to EWSR1::FLI1.

Our functional experiments demonstrate that *GLRX3* silencing strongly inhibits tumor growth in vivo, which is associated with drastic dysregulation of the oxidative stress and iron homeostasis. However, whether these phenotypes are mechanistically connected and how GLRX3 mediates them needs to be explored in subsequent studies, especially regarding future therapeutic exploitation.

Although we cannot extrapolate with certainty from other cancer entities to EwS, it should be noted that prior studies in nasopharyngeal carcinoma cell lines (HONE1, CNE2), cervical cancer cells (HeLa) and immortalized T lymphocytes (Jurkat) have shown that GLRX3 attenuates oxidative stress levels^22,63,64^. Similarly, previous studies have shown that *GLRX3* silencing may result in dysregulation of cellular iron homeostasis by decreased ISC activity and accumulation of iron^25,65^, which is why it is tempting to speculate that similar mechanisms might be operative in EwS, too. The preferential dependency on GLRX3 suggested that this protein could constitute an attractive target structure for treatment of EwS patients. Since there are no specific GLRX3 inhibitors available to date, we explored the possibility of using this protein as a predictive biomarker for already available drugs that have increased efficacy in either GLRX3-high or -low conditions. In line with the observed pro-proliferative effect of GLRX3 (**Fig. 2**), our experiments indicated that GLRX3-high EwS cells are more susceptible toward targeted inhibition of CDK4/6 (**Fig. 4**). These data are in keeping with previous reports from us and others suggesting a high sensitivity of EwS cells for the CDK4/6 inhibition^52–54^. In contrast to these promising preclinical signals, a recent phase II clinical trial employing palbociclib in relapsed EwS patients did not detect a remarkable overall response. Yet, it is worth noting that some patients did show a moderate response and that the trial was conducted without biomarker guidance^51^. Thus, it remains to be determined whether CDK4/6 inhibition could yield better outcomes when preselecting/stratifying for GLRX3-high patients, and by possibly also including patients at a less advanced disease stage.

In analogy to our approach to target GLRX3-high cells, we demonstrated that GLRX3-low EwS cells are more sensitive toward apoptosis regulator Bcl-2 and Bcl-xL inhibition with navitoclax (**Fig. 4**), which is being under clinical investigation for mainly hematologic malignancies (*NCT00445198*, *NCT01989585*, *NCT06210750*, *NCT02079740*, *NCT05740449*). Since – similar to trials testing CDK4/6 inhibitors – most previous clinical trials testing navitoclax have been conducted without biomarker-guided patient stratification, it is intriguing to see whether GLRX3 expression could serve as a therapeutic biomarker to predict a selective antitumor effect by navitoclax in EwS. In addition to navitoclax, we further uncovered erastin as another drug selectively efficacious for GLRX3-low EwS cells (**Fig. 4**). Erastin inhibits cystine-glutamate antiporter and thereby depletes a major cellular antioxidant glutathione, rendering cells more vulnerable to ferroptotic cells death^57,59^. Our observation that GLRX3 silencing remarkably increased oxidative stress level in EwS (**Fig. 3**) suggests that the augmentation of oxidative stress and disruption of iron homeostasis through GLRX3 silencing sensitizes EwS cells toward ferroptotic cell death, which is eventually mediated by cell membrane lipid peroxidation^55,56^. However, due to its poor bioavailability, erastin is currently limited to experimental application. Thus, it will be of interest to test whether other drugs reported to promote ferroptotic cell death (e.g. sorafenib, sulfasalazine)^57^ could also demonstrate selectivity for GLRX3-low EwS cells. Yet, to the best of our knowledge, there are to date no alterative and highly specific ferroptosis inducers available that show promising pharmacological features that could enable a rapid clinical translation^57^.

In synopsis, our results exemplify how the interplay of the evolutionarily conserved oxidative stress regulator GLRX3 with dominant EWSR1::FLI1 fusions can promote malignancy, which may also provide opportunities for predictive diagnostics and personalized therapy of EwS patients.

## MATERIALS AND METHODS

### Provenience of cell lines and cell culture conditions

The human EwS cell line A-673 (RRID:CVCL 0080) was purchased from American Type Culture Collection (ATCC). Human EwS cell line MHH-ES-1 (RRID:CVCL 1411) and human HEK293T cells were provided by the German Collection of Microorganisms and Cell cultures (DSMZ). Human EwS cell line EW-22 was kindly provided by O. Delattre (Institut Curie, Paris). All cell lines were maintained under standardized conditions at 37 °C in a humidified atmosphere containing 5% CO_2_ in RPMI 1640 medium (Sigma-Aldrich, Germany) supplemented with stable glutamine (Biochrom, Germany), 10% tetracycline-free fetal calf serum (FCS) (Sigma-Aldrich), and penicillin (Biochrom) at a concentration of 100 U/ml, and streptomycin (Biochrom) at a concentration of 100 µg/ml. Regular screening for mycoplasma infection was performed using nested PCR. The purity and authenticity of the cell lines were further confirmed by STR-profiling.

### Nucleotide extraction, reverse transcription, and quantitative real-time PCR (qRT-PCR)

Total RNA extraction was carried out utilizing the NucleoSpin RNA kit (Macherey-Nagel, Germany) to isolate the RNA samples. Subsequently, 1 µg of total RNA was subjected to reverse transcription using the High-Capacity cDNA Reverse Transcription Kit (Applied Biosystems, USA). For quantitative real-time polymerase chain reaction (qRT-PCR), SYBR green Mastermix (Applied Biosystems) was employed, along with diluted cDNA (1:10) and 0.5 µM forward and reverse primers (total reaction volume 16 µl). The qRT-PCR reactions were performed on a BioRad CFX Connect instrument, and the resulting data were analyzed using the BioRad CFX Manager 3.1 software. To determine gene expression levels, the 2^−(ΔΔCt)^ method was utilized, normalizing the values relative to the housekeeping gene *RPLP0* as an internal control. Oligonucleotides required for the qRT-PCR experiments were purchased from MWG Eurofins Genomics (Germany) and are listed in **Supp. Table 4**. The thermal conditions for qRT-PCR were as follows: initial heat activation at 95 °C for 2 min, followed by DNA denaturation at 95 °C for 10 s, and subsequent annealing and elongation at 60 °C for 20 s (50 cycles). Finally, a final denaturation step was performed at 95 °C for 30 s.

### Generation of doxycycline (Dox)-inducible shRNA constructs

The human EwS cell lines A-673, EW-22, and MHH-ES-1 were transduced with lentiviral Tet-pLKO-puro all-in-one vector system (plasmid #21915, Addgene) containing a puromycin-resistance cassette, and a tet-responsive element for Dox-inducible expression of shRNAs against *GLRX3* (shGLRX3_1 or shGLRX3 _2) or a non-targeting control shRNA (shCtrl). Sequences of the used shRNAs are listed in **Supp. Table 4**. Dox-inducible vectors were generated according to a publicly available protocol^66^, using In-Fusion HD Cloning Kit (Takara Bio, USA). Vectors were amplified in Stellar Competent Cells (Takara Bio) and respective integrated shRNA was verified by Sanger sequencing. The used sequencing primer is listed in **Supp. Table 4**. Lentiviral particles were generated in HEK293T cells. Virus-containing supernatant was collected to infect the human EwS cell lines. Successfully transduced cells were selected with 1 µg/ml puromycin (InVivoGen, France). The shRNA expression for *GLRX3* knockdown or expression of a negative control shRNA in EwS cells was achieved by adding 0.1 µg/ml Dox every 48 h to the medium. Generated cell lines were designated as A-673/TR/shCtr, A-673/TR/shGLRX3_1, A-673/TR/shGLRX3_2, MHH-ES-1/TR/shCtr, MHH-ES-1/TR/shGLRX3_1, MHH-ES-1/TR/shGLRX3_2, EW-22/TR/shCtr, EW-22/TR/shGLRX3_1, and EW-22/TR/shGLRX3_2.

### Western blot

EwS harboring inducible shRNAs were treated for 96 h with Dox to induce *GLRX3* knockdown. Whole cellular protein was extracted with RIPA buffer (Tris buffer pH 7.5 25mM, NaCl 150mM, Tergitol 15-s-9 1%, desoxycholic acid-Na-salt 1%, SDS 0.1%) (Serva electrophoresis, Germany) containing protease inhibitor cocktail and phosphatase inhibitor cocktail (Roche, Switzerland). Western blots were performed following routine protocols^67^. Detection of specific protein bands was accomplished using a rabbit monoclonal anti-GLRX3 antibody (1:1,000, HPA028941, Sigma-Aldrich), ferritin (1:1,000, AB69090, Abcam, UK), TfR1 (1:1,000, 136800, Invitrogen/Life Tech) and a mouse monoclonal anti-ß-actin antibody (1:5,000, A-5441, Sigma-Aldrich and 1:1,000, A1978, Sigma-Aldrich). Afterwards, the nitrocellulose membranes (GE Healthcare BioSciences, Germany) were subjected to a secondary incubation with an Anti-rabbit IgG horseradish peroxidase coupled antibody (1:2,000, sc-516102, Santa Cruz Biotechnology, Germany) and an anti-mouse IgG horseradish peroxidase coupled antibody (1:2,000, sc-2357, Santa Cruz Biotechnology). Protein bands were visualized using a chemiluminescence HRP substrate (Merck, Germany).

### Proliferation assays

Depending on the cell line, 3–5 **×** 10^4^ EwS cells containing either a Dox-inducible non-targeting control shRNA or *GLRX3*-targeting specific shRNAs were seeded in wells of a 6-well plate in 2 ml of growth medium. The cells were treated either with or without Dox (0.1 µg/ml every 48 h; Sigma-Aldrich) for 96 h. Subsequently, the cells from each treatment group, including the supernatant, were collected and subjected to Trypan blue staining (Sigma-Aldrich). Viable and non-viable cells were then quantified using a standardized hemocytometer (C-chip, NanoEnTek, South Korea).

### Clonogenic growth assays (Colony forming assays)

EwS cell lines containing either a Dox-inducible non-targeting control shRNA or *GLRX3*-targeting specific shRNAs were seeded in triplicate wells of a 6-well plate at a density of 3,000 cells (A-673), 3,500 (MHH-ES-1) or 4,000 cells (EW-22) per well in 2 ml of growth medium. Cells were grown with/without Dox (0.1 µg/ml; Sigma-Aldrich) for 11 days. Following that, the colonies were treated with Crystal Violet solution (Sigma-Aldrich), and subsequently, the number and size of the colonies were quantified using the ImageJ Plugin Colony area.

### Cell cycle analysis

To conduct cell cycle analysis, A-673, EW-22, and MHH-ES-1 cells containing a Dox-inducible shRNA targeting *GLRX3* were synchronized by a double thymidine (T1850, Sigma-Aldrich) block/release^68^. In brief, cells were initially blocked in the G1/S phase by treating them with 1 mM thymidine for 18 h at 37 °C. Subsequently, the cells were released into the S phase by washing them three times with pre-warmed serum-free media. Afterwards, fresh complete medium was added, and the cells were incubated for 10 h at 37 °C. To bring the cells back to the G1/S boundary, a second round of thymidine at a final concentration of 1 mM was introduced, and the cells were cultured for an additional 18 h at 37 °C. At this point, the cells were in the G1/S boundary state. Following three washes with pre-warmed serum-free media, the cells were seeded at a density of 5 × 10^5^ cells per T25 flask in fresh medium with/without Dox (0.1 µg/ml). The Dox treatment was renewed 48 h after seeding. After 96 h the cells were fixed with ice-cold 70% ethanol, treated with 100 µg/ml RNAse (ThermoFisher, USA), and stained with 50 µg/ml propidium iodide (PI) (Sigma-Aldrich). Samples were assayed by BD FACSCanto II (BD Biosciences, USA).

### Apoptosis analysis via Annexin V/PI staining

For these assays, EwS cells containing a Dox-inducible shRNA against *GLRX3* were seeded in triplicate wells of 6-well plates at a density of 3–5 × 10^3^ cells per well in 2 ml growth medium and treated with 0.1 µg/ml Dox every 48 h. After 96 h, the cells were washed with phosphate buffered saline (PBS) and resuspended in 1× Annexin V buffer (BD Biosciences) containing 5 µl of Annexin V and 5 µl of PI solution for an additional 15 min. Analysis of Annexin V/PI positivity was conducted using the BD FACSCanto II (BD Biosciences) by evaluating at least 1 × 10^4^ events.

### Oxidative stress detection via DCF-DA fluorescence

To assess oxidative stress, EwS cells containing a Dox-inducible shRNA against *GLRX3* were seeded at a density of 3–5 × 10^3^ cells per 2 ml per 6-well plate. These cells were directly treated with Dox for 96 h. On the day of analysis, the cells were incubated in their respective medium containing 2.5 µM DCF-DA (ThermoFisher) for 30 min at 37 °C. Subsequently, the cells were harvested and suspended in PBS for flow cytometry analysis using FACSCanto II (BD Biosciences).

### Drug screening assays (3D)

For the drug screening experiments, a methodology previously described^69^ was followed. 384-well round bottom ultralow attachment spheroid microplates (# 3830, Corning, USA) were utilized to facilitate the formation of three-dimensional spheroids. In summary, a drug library comprising 87 compounds was assessed, primarily consisting of approved drugs or those in clinical trials. The library encompassed standard chemotherapeutic drugs, epigenetic modifiers, metabolic modifiers, kinase inhibitors, apoptotic modulators, and other compounds^69^. Drug plates were obtained as pre-prepared assay plates from the FIMM High Throughput Biomedicine Unit (Institute for Molecular Medicine Finland HiLIFE, University of Helsinki, Finland) and stored in a controlled environment (San Francisco StoragePod, Roylan Developments Ltd, Fetcham Leatherhead, UK) at room temperature, protected from oxygen and moisture. Each set of drug plates consisted of three individual plates. The concentration range of each drug spanned five orders of magnitude, with each condition tested in duplicate. Control wells contained 100 µM benzethonium chloride (BztCl), 250 nM staurosporine (STS), and 0.1% dimethyl sulfoxide (DMSO), representing maximum, intermediate, and minimum effect controls, respectively. An STS concentration range of 0.1 to 1,000 nM was included as a technical control, with four replicates per plate.

A volume of 25 µl of single-cell suspension was dispensed into each well using an 8-channel electronic Picus^®^ pipette (10–300 µl, #735361, Sartorius, Germany). To evaluate cell viability, bulk ATP quantitation was performed 72 h after treatment using CellTiter-Glo^®^ 2.0 (CTG; #G9243, Promega, Germany), following the manufacturer’s instructions. This enabled the determination of the relative number of metabolically active cells per well. Prior to conducting the drug screen, we determined the suitable cell number to be seeded per well by assessing the relationship between the number of cells and the luminescence signal. We tested 200, 500, 1,000, and 2,000 cells in the absence of drug treatment. Based on this assessment, 250 cells per well were selected for A-673/TR/shGLRX3 Dox (–) and 350 cells per well for A-673/TR/shGLRX3 Dox (+) for the subsequent screening experiment. To calculate the drug effects, we employed drug sensitivity scores (DSSasym) using the web-based drug analysis pipeline iTReX61 (https://itrex.kitz-heidelberg.de/iTReX/)^40^. The differential DSS (ΔDSS) was determined by calculating the difference between the DSSasym values of GLRX3-high and -low cells for each drug.

### Drug-response assays in vitro

A-673 and EW-22 EwS cells were plated in a 96-well plate with a density of 2–4.5 × 10^3^ cells per well. In the case of cells containing Dox-inducible constructs, they were either treated with Dox (0.1 µg/ml; Sigma-Aldrich) or left untreated. After 48 h of seeding or pre-incubation with Dox, respectively, a series of diluted concentrations of erastin [0.125 µM – 16 µM], navitoclax [0.25 µM – 32 µM], palbociclib [0.78 µM – 100 µM] and ribociclib [2 µM – 250 µM] were added to the wells. Each well contained less than 0.5% DMSO (Sigma-Aldrich) as a solvent. Cells treated only with 0.5% DMSO were used as a control. Following 72 h of treatment, the plates were analyzed using a GloMax^®^ Explorer Multimode Microplate Reader after incubating with Resazurin (16 µg/ml; Sigma-Aldrich) for 6–8 h.

### *In vivo* experiments in mice

NOD/Scid/gamma (NSG) mice aged 10–12 weeks were housed in individually ventilated cages (IVC) under specific pathogen-free (SPF) conditions with controlled dark/light cycles, temperature, and humidity. EwS cells, either containing a specific shRNA or a non-targeting control, were suspended in a 1:1 mix of PBS and Geltrex Basement Membrane Mix (ThermoFisher). Thereafter, 2.5 × 10^6^ cells were injected subcutaneously into the right flank of the NSG mice (Charles River Laboratories, USA). Tumor diameters were measured every second day using a caliper, and tumor volume was calculated based on the formula *V= (a × b^2^)/2*, where *a* represents the largest diameter and *b* represents the smallest diameter of the tumor. Once the tumors were palpable, the mice were randomized into two groups. The treatment group received 2 mg/ml Dox (BelaDox) dissolved in drinking water containing 5% sucrose (Sigma-Aldrich), to induce an in vivo knockdown, while the control group only received water containing 5% sucrose. Tumor growth was monitored regularly until the control group’s tumors nearly reached a size of nearly 15 mm in one dimension. At this point, all mice in the experiment were euthanized by cervical dislocation.

Various humane endpoints, including ulcerated tumors, body weight loss, body posture changes, digestive abnormalities, breathing difficulties, dehydration, abdominal distention, body condition scores, apathy, and self-isolation, were considered to determine the welfare of the animals throughout the experiment. All mouse experiments were conducted in compliance with ethical regulations and received approval from local authorities.

### Gene expression microarrays

To evaluate the influence of GLRX3 on gene expression in EwS, a microarray analysis was conducted. A-673/TR/shGLRX3 and EW-22/TR/shGLRX3 EwS cells were cultured in T25 flasks and treated with Dox (0.1 µg/ml, Sigma-Aldrich) for 96 h, with Dox refreshment after 48 h. Similarly, we cultured EW-22/TR/shCtr cells. Total RNA was extracted using the Nucleospin II kit (Macherey-Nagel), and the transcriptomes were profiled at IMGM laboratories (Martinsried, Germany) using Human Affymetrix Clariom D microarrays. RNA quality was assessed with a Bioanalyzer, and samples with RNA integrity numbers (RIN) greater than 9 were selected for hybridization. The resulting data were either combined with previous data for A-673/TR/shCtr (GSE166415)^70^ or with A-673/TR/shEF1 and EW-22/TR/shEF1 cells (GSE176190)^36^. Data were jointly quantile normalized using Transcriptome Analysis Console (v4.0; ThermoFisher Scientific) with the Signal Space Transformation – Robust Multi-array Average (SST-RMA) algorithm. Gene-level annotation was performed using the Affymetrix library for Clariom D Array (version 2, human). DEGs with consistent and significant fold changes (FCs) across shRNAs and cell lines were identified. Log2 FCs for specific shRNAs in each cell line were calculated by subtracting the log2 FCs of control samples from those of the corresponding shRNA samples. Additionally, for differentially expressed genes across shRNAs and cell lines, a mean log2 FC was calculated for each cell line/construct and summarized and normalized to the control shRNA. Statistical significance levels of DEG-overlaps between conditions were calculated using a Chi-square test.

### Fast gene-set enrichment analysis (fGSEA) and single-sample GSEA (ssGSEA)

For the identification of enriched gene sets, genes were sorted based on their FC in expression between the two groups, namely Dox (−/+) group. A pre-ranked GSEA was performed using the fgsea package (version 1.14.0) in R (1,000 permutations). The analysis utilized the biological process Gene Ontology (GO) definitions from MSigDB (version 7.5.1) with the symbol-based gene set files c2.cp.reactome and c5.go.bp (symbols.gmt). By utilizing the Affymetrix gene expression dataset consisting of 196 EwS patients (GSE63157, GSE34620, GSE12102, GSE17618 and unpublished data), the enrichment of gene sets co-regulated with GLRX3 was determined. This was achieved by ranking the Pearson’s correlation coefficient of each gene’s expression with the expression of GLRX3, followed by conducting a pre-ranked fGSEA. To assess the correlation between GLRX3 expression and gene enrichment scores in EwS patient tumors, we applied single sample GSEA (ssGSEA)^71,72^ on the EwS patient cohort of 196 tumor samples mentioned above.

### Histology

Routine protocols were followed for hematoxylin and eosin (HE)-staining of EwS xenografts. The quantification of mitoses in HE-stained slides of EwS xenografts was performed by two blinded observers, who assessed 5 high-power fields per sample. The average number of mitoses per sample was calculated based on a total of 10 high-power fields counted by both observers.

### DepMap dependency screen analysis

The ‘DepMap public 23Q4+Score, Chronos’ data set was used to investigate gene dependency effects in EwS cell lines and those of other cancer entities. EwS specific dependencies were calculated by performing a two-tailed t-test for the difference in gene dependency effect between EwS cell lines (*n*=23) and other cancer entities (*n*=1,077). Statistical significance was calculated using the Benjamini & Hochberg^73^ method resulting in adjusted *P*-values.

### Statistics and software

Statistical analysis was conducted using GraphPad PRISM 10 (GraphPad Software Inc., CA, USA) on the raw data. If not otherwise specified in the figure legends, a two-tailed Mann-Whitney test was used to compare two groups in functional in vitro experiments. Unless stated otherwise in the figure legends, data were presented as dot plots with mean values represented by horizontal bars and standard error of the mean (SEM) represented by whiskers. The sample size for all in vitro experiments was determined empirically, ensuring a minimum of three biological replicates. Pearson’s correlation coefficients were calculated using either Microsoft Excel or GraphPad PRISM 10 (GraphPad Software Inc., CA, USA). For in vivo experiments, sample size was predetermined through power calculations, considering *β* = 0.8 and *α*=0.05, based on preliminary data and adhering to the principles of the 3R system (replacement, reduction, refinement).

## Data availability

In this study, we used DNA microarray data based on Affymetrix Clariom D arrays deposited at the Gene Expression Omnibus (GEO) for A-673/TR/shEF1 and EW-22/TR/shEF1 cells (GSE176190) as well as A-673/TR/shCtr cells (GSE166415). In addition, we generated new DNA microarray data on the same array platform that is also available via the GEO (GSE264510).

## ACKNOWLEGEMENTS

We thank Stefanie Kutschmann, Felina Zahnow, Nadine Gmelin, and Aileen Friedenauer for excellent technical assistance. Support by the DKFZ Light Microscopy Facility is gratefully acknowledged.

## AUTHOR CONTRIBUTIONS

E.V., A.C.E., A.R., D.O., C.M.F., F.H.G., M.S., J.L., M.Z., A.K.C., and F.F. performed functional in vitro and in vivo experiments, as well as bioinformatic and histological analyses. J.A. provided cell line models and microarray data. H.P., O.W., and I.O., performed drug screens. F.C.A., R.I., A.B. helped in coordination and performance of in vivo experiments. C.M., R.Q., M.M.M., A.D., and S.v.K. coordinated and carried out experiments regarding iron metabolism and cell death. T.G.P.G. and S.O. designed and supervised the study, provided biological and technical guidance, analyzed and interpreted all data, and provided laboratory infrastructure. All authors read and approved the final manuscript.

---

**Supp. Fig. 1.**
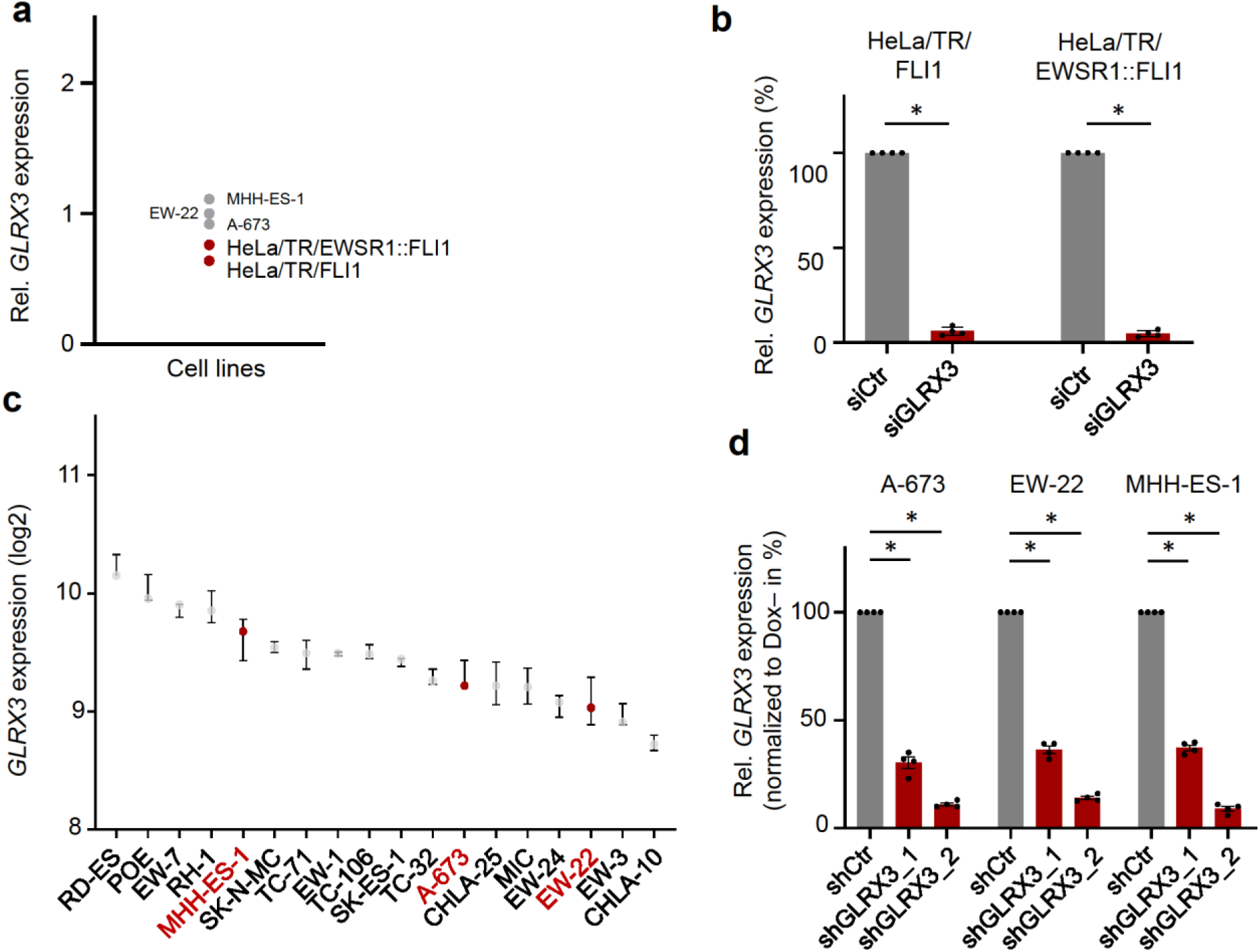
*GLXR3* expression and its siRNA/shRNA-mediated knockdown in EwS and HeLa cells. **a)** Relative *GLRX3* expression in HeLa/TR/EWSR1::FLI1 and HeLa/TR/FLI1 cells compared to A-673, EW-22 and MHH-ES-1. **b)** Relative *GLRX3* expression in HeLa/TR/EWSR1::FLI1 and TR/FLI1 cells after 72h of siRNA transfection. *n*=4 biologically independent replicates. Horizontal bars represent means, and whiskers represent the SEM. **c)** *GLRX3* expression levels (Affymetrix microarray) in 18 EwS cell lines. A-673, EW-22, and MHH-ES-1 highlighted in red. Horizontal bars represent means, and whiskers represent the SEM. *n*=3 biologically independent replicates **d)** Relative *GLRX3* expression in A-673, EW-22 and MHH-ES-1 96 h after shRNA-mediated knockdown. Horizontal bars represent means, *n*=4 biologically replicates. *P*-values determined by two-sided Mann-Whitney test. \**P* < 0.05

**Suppl. Fig. 2.**
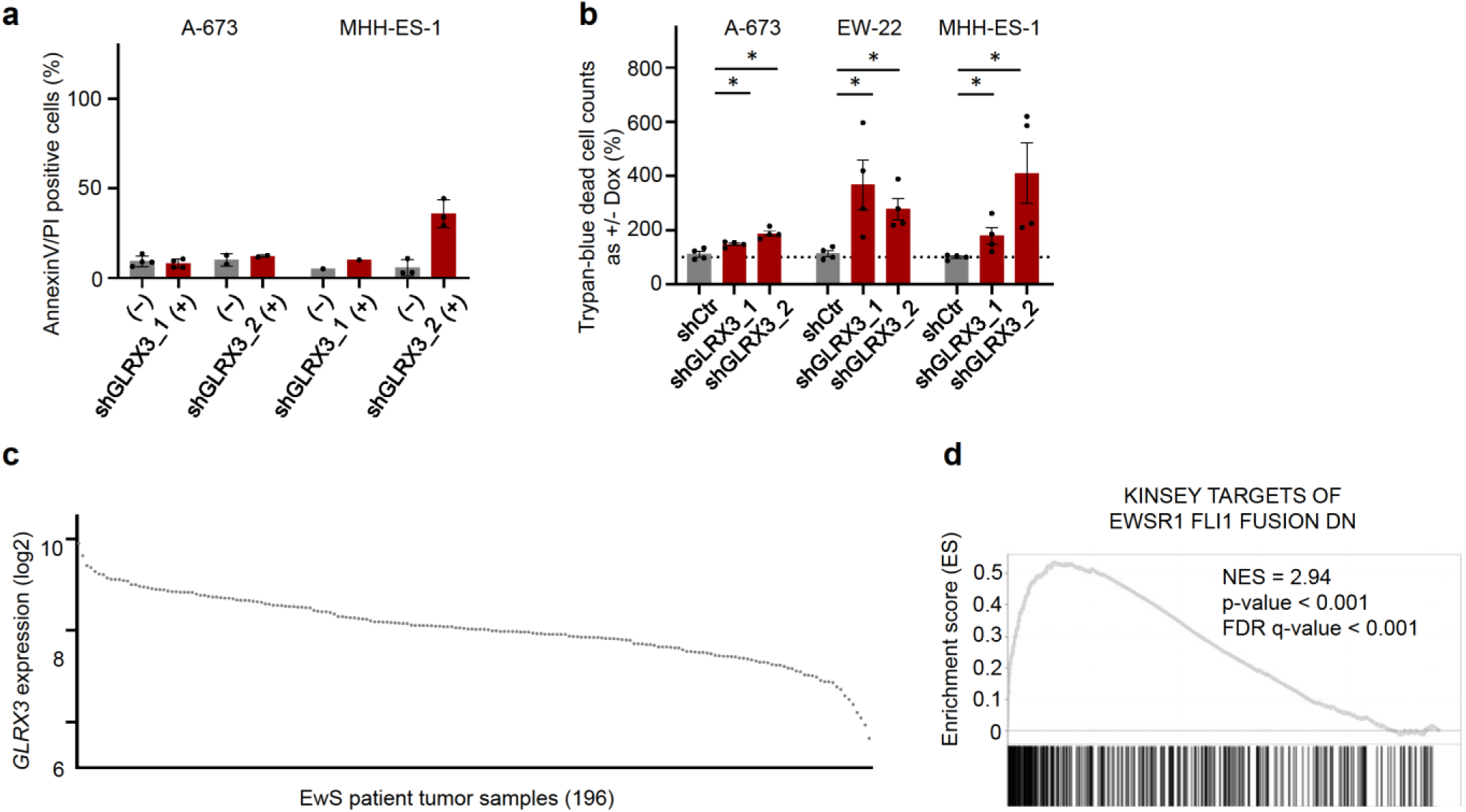
GLRX3 silencing effects on cell viability, and GLRX3 expression in EwS tumor samples. **a)** Percentage of AnnexinV/PI positive cells 96 h after Dox-treatment. Horizontal bars represent means, error bars represent SEM, *n*≥1 biologically independent replicate(s). **b)** Percentage of dead cell counts assessed by Trypan-blue staining 96 h after Dox-treatment. Horizontal bars represent means, error bars represent SEM, *n*=4 biologically independent replicates. \**P* < 0.05. **c)** *GLRX3* expression levels in 196 EwS tumors. **d)** Most enriched pathway after *GLRX3* silencing assessed by GSEA in c2.cgp gene set is the downregulation of *EWSR1::FLI* fusion targets.

